# Characterization of raloxifene as potential pharmacological agent against SARS-CoV-2 and its variants

**DOI:** 10.1101/2021.10.22.465294

**Authors:** Daniela Iaconis, Carmine Talarico, Candida Manelfi, Maria Candida Cesta, Mara Zippoli, Francesca Caccuri, Giulia Matusali, Licia Bordi, Laura Scorzolini, Enrico Bucci, Arnaldo Caruso, Emanuele Nicastri, Marcello Allegretti, Andrea Rosario Beccari

## Abstract

The new coronavirus that emerged, called SARS-CoV-2, is the causative agent of the COVID-19 pandemic. The identification of potential drug candidates that can rapidly enter clinical trials for the prevention and treatment of COVID-19 is an urgent need, despite the recent introduction of several new vaccines for the prevention and protection of this infectious disease, which in many cases becomes severe. Drug repurposing (DR), a process for studying existing pharmaceutical products for new therapeutic indications, represents one of the most effective potential strategies employed to increase the success rate in the development of new drug therapies. We identified raloxifene, a known Selective Estrogen Receptor Modulator (SERM), as a potential pharmacological agent for the treatment of COVID-19 patients. Following a virtual screening campaign on the most relevant viral protein targets, in this work we report the results of the first pharmacological characterization of raloxifene in relevant cellular models of COVID-19 infection. The results obtained on all the most common viral variants originating in Europe, United Kingdom, Brazil, South Africa and India, currently in circulation, are also reported, confirming the efficacy of raloxifene and, consequently, the relevance of the proposed approach.

Taken together, all the information gathered supports the clinical development of raloxifene and confirms that the drug can be proposed as a viable new option to fight the pandemic in at least some patient populations. The results obtained so far have paved the way for a first clinical study to test the safety and efficacy of raloxifene, just concluded in patients with mild to moderate COVID-19.

## Introduction

Coronaviruses are the causative agent of multiple respiratory and intestinal infection in humans and animals [1-3]. Unlike most other human coronaviruses, which only rarely cause severe disease and death [4], SARS-CoV-2 is able to cause severe acute respiratory illness, multi-organ failure and death, sharing common pathogenetic mechanisms with SARS-CoV and MERS-CoV betacoronavirus [5]. Common symptoms of COVID-19 include fever, sore throat, fatigue, cough, shortness of breath and dyspnea that may eventually progress towards acute respiratory distress syndrome, with the involvement of other systems/organs (e.g., heart, liver and kidneys) [6, 7] and up to the death in the most critical cases. About 80% of patients have mild to moderate disease, 14% have severe disease, and 6% become critical (namely, they develop respiratory failure, septic shock, and/or multiple organ dysfunction/failure) (www.ecdc.europa.eu – fifth update, 2 March 2020). As of September 6^th,^ 2021, SARS-CoV-2 infection led to more than 4,5 million deaths worldwide (https://covid19.who.int/). To date, notwithstanding the advent of vaccine programs and constant social distancing interventions, it is believed that the virus is likely or very likely to become endemic [8, 9]. In addition, the emerging of SARS-CoV-2 variants raises great concern for vaccine efficacy, reinfection events and increased transmissibility and disease severity. As the virus started to spread around the world, a mutated spike SARS-CoV-2 variant (D614G) emerged and was associated with increased infectivity, becoming the predominant variant in Europe and worldwide without any increase in disease severity [10-13]. In recent months, other variants were defined as “variants of concern” (VOC). The most relevant also from a clinical point of view are: B.1.1.7 (UK), B.1.351 (South African), B.1.1.28 (Brazilian P.1), B.1.427 and B.1.429 (Californian, also named CAL.20C), characterized by increased transmissibility, immune evasion and higher virulence [14-20]. As of May 11^th^, 2021, the so called “Indian” Delta variant (Delta B.1.617.2) was added to the WHO list of VOC; this variant seems to be able to escape adaptive immunity induced by prior wild type infection roughly half of the time and to be more infectious (around 60%) than wild type SARS-CoV-2 [21]. The growing relevance of the rapidly emerging SARS-CoV-2 variants deserves further investigations, and new impetus will have to be given to research to increase the availability of broad-spectrum drugs or vaccines for long-term prevention, treatment and control of COVID-19, with the final goal to find new interventions and cures to complement vaccine programs. To identify potential therapeutic targets, one of the main studied mechanisms is the virus entry machinery, and several preclinical and clinical trials are ongoing to find new inhibitors of clinical relevance [22]. The entry machinery involves two key host proteins: the angiotensin-converting enzyme 2 (ACE2) and the cell surface transmembrane protease serine 2 (TMPRSS2) [23, 24]. In addition, also Neuropilin1 (NRP-1) has been recognized as an important receptor whose inhibition reduces SARS-CoV-2 entry and infectivity [25, 26].

Moreover, recent evidence shows that Nuclear Receptors (NRs), and in particular the sex hormone receptors, like estrogen and androgen receptors, could be involved in the outcome of COVID-19. These receptors regulate the viral entry protein expression and activity [27-29]. Additionally, a protective effect of estrogens in the progression of COVID-19 infection has been associated with their role in regulation of innate and adaptive immune responses, as well as in the control of the cytokine storm [30-34], whereas activation of androgen receptors seems to correlate with the worse COVID-19 clinical outcome observed in men compared to women [29, 35-37].

Recently, several molecules with potential efficacy against SARS-CoV-2 were selected from an extensive virtual screening campaign based on the EXaSCale smArt pLatform Against paThogEns [38], a powerful tool for repurposing of drugs and compounds in new indications [39-42] for immediate response and quick identification of effective treatments, useful during pandemic situations. So, in the context of the Horizon 2020 project EXSCALATE4CoV, raloxifene, a well-known SERM (Selective Estrogen Receptor Modulator) [43-47] was selected through an integrated approach of drug repurposing and *in silico* screening on SARS-CoV-2 target proteins, an approach that, combined with the scientific rationale and literature evidence that support a potential antiviral and protective action of SERMs in COVID-19, led the molecule to be selected as clinical candidate for studies in mild to moderate COVID-19 patients [38].

Raloxifene is a drug registered in Europe and US for the treatment and prevention of osteoporosis in postmenopausal women, and for the reduction of the risk of invasive breast cancer in postmenopausal women [48, 49]. It is known to act as an agonist in the bone, liver and cardiovascular system and as antagonist in human breast and uterine tissues [50-52], and tissue specificity is relevant for its use in postmenopausal osteoporosis and prevention of breast cancer without increase of risk of endometrial cancer, differently from the behavior of other SERMs like tamoxifen [53, 54]. The drug has also been studied in men for uses such as for treatment of schizophrenia, prostate cancer and osteoporosis [55-57]. Recently, raloxifene has been also characterized in viral infections. It was found active against Ebola virus [58, 59], Hepatitis C virus [60, 61], Hepatitis B virus [62], and Zika virus [63]. Further, it showed efficacy in human female cells from nasal epithelium, against the Influenza Virus A [64], and as adjuvant antiviral treatment of chronic hepatitis C (CHC) in postmenopausal women [65]. These observations, together with raloxifene activity on the Estrogen Receptor (ER) pathways, highlight a possible relationship between clinical outcome and sex and age of patients with viral infections.

In this paper, we report for the first time a full characterization of the antiviral activity of raloxifene in two different well-established cellular contexts (Vero E6 and Calu-3) and we tested the potential influence of the most common COVID-19 variants on raloxifene biological activity. Raloxifene *in vitro* activity was high and consistent in the different cell lines tested, preserved in all main VOCs of clinical relevance.

Repurposing and *in silico*/experimental synergy are powerful useful approaches in case of pandemic infections by viruses and other pathogens, where an immediate response and the swift identification of effective treatments are of paramount importance. Taken together, the collected evidence confirms the potential of raloxifene as a promising agent with the potential to control COVID-19 infection with pleiotropic mechanisms supporting the rationale for the ongoing clinical investigation (study RLX0120, EudraCT Nr: 2020-003936-25) for the treatment of mild to moderate COVID-19 patients.

## Materials and methods

### System Biology Screening

First, we isolated 12 genes identified by The Host Genetic Initiative (https://www.covid19hg.org/results/r3/) as relevant for the infection and present in the data release number 3 (July 2020), namely: ANKRD32, CDRT4, PSMD13, ERO1L, LZTFL1, XCR1, FYCO1, IFNAR2, CXCR6, CCR9, AP000295.9, AK5. Based on a lookup of previous GWAS results in the GWAS ATLAS database (a database of publicly available GWAS summary statistics), these genes are considered primarily implicated in immunological phenotypes. Then, we looked at the human-SARS-CoV-2 interactome network as published [66], and extracted all the human genes included in the set. A list combining the two dataset was used as seeding for a BioGrid search by mean of Cytoscape v.3.8.0; the resulting enriched functional network connected those human proteins, known to directly bind SARS-CoV-2 proteins, with the human gene products involved in the host pathology. Subsequently, we screened 8721 Scopus-derived documents, referred to raloxifene, for the presence of at least one of the proteins/genes included in the Cytoscape-generated network; this allowed to isolate 600 papers, which were manually examined and annotated for enriched human gene ontology according to BiNGO v.3.5.0.

### Cells

African green monkey kidney Vero E6 cell line was obtained from American Type Culture Collection (ATCC, Manassas, VA, USA) and maintained in Dulbecco’s Modified Eagle Medium (DMEM; Gibco, Thermo-Fisher, Waltham, MA, USA) supplemented with 10% fetal bovine serum (FBS; Gibco, Thermo-Fisher) at 37°C in a humidified atmosphere of 5% CO_2_.

Calu-3 (human, Caucasian, lung, adenocarcinoma) cell line was obtained from ATCC and maintained in Minimum Essential Medium (MEM; Gibco, Thermo-Fisher) supplemented with 10% fetal bovine serum (FBS; Gibco, Thermo-Fisher) at 37°C in a humidified atmosphere of 5% CO_2_.

### Virus

Different SARS-CoV-2 variants isolated from COVID-19 patients’ respiratory samples were used. The identity of each variant was verified by metagenomic sequencing. Genomic data of SARS-CoV-2, belonging to the B.1 lineage, are available at EBI (under study accession number: PRJEB38101) [67, 68].

Below the list of the viral strains used to assess the activity of raloxifene against viral variants:

- Human 2019-nCoV strain 2019-nCoV/Italy-INMI1, clade V (Ref-SKU: 008V-03893, EVAg portal), and isolated in January 2020 from a chinese patient (control infection) (named Wuhan)
- SARS-CoV-2 isolate SARS-CoV-2/Human/ITA/PAVIA10734/2020, clade G, D614G (S) (Ref-SKU: 008V-04005, EVAg portal), named D614G, isolated in Lombardy in February 2020
- SARS-CoV-2 isolate hCoV-19/Italy/LAZ-INMI-82isl/2020, clade GV, A222V, D614G (S) (Ref-SKU: 008V-04048, EVAg portal), named GV and representig the dominant strain circulating in Europe from April to December 2020.
- SARS-CoV-2 variant VOC 202012/01, isolate hCoV-19/Italy/CAM-INMI-118isl/2020, clade GR, Δ69-70, Δ144, N501Y, A570D, D614G, P681H, T716I (S) (Ref-SKU: 008V-04050, EVAg portal), named VOC B.1.1.7 and representing the variant of major concern from UK
- SARS-CoV-2 variant GR/501Y.V3, isolate hCoV-19/Italy/LAZ-INMI-216isl/2021, clade GR, PANGO lineage P.1, K417T, E484K, N501Y (S) (Ref-SKU: 008V-04101, EVAg portal), named VOC P1 representing the variant of major concern from Brazil
- SARS-CoV-2 variant VOC SA/B.1.351, obtained by GHSAG (Public Health England), named B.1.351 and representing the variant of major concern from South Africa
- SARS-CoV-2 variant VOC G Delta /B1.617.2 isolate hCoV-19/Italy/LAZ-INMI-648/2021 (EPI_ISL_2000624), named VOC B.1.617.2 representing the variant of major concern from India

All the infection experiments were performed in a biosafety level-3 (BLS-3) laboratory at a multiplicity of infection (MOI) of 0.05.

### Cell viability studies of raloxifene

Cells were seeded into 24-well plates (2.5×10^4^ cells/well) in DMEM supplemented with 10% FBS, and treated with different doses of raloxifene (1.25, 2.5, 5, 10, 15, 20, 25 and 30 µM) at 37°C for 48 h. Cell viability was estimated by measuring the ATP levels using CellTiter-Glo (Promega, Madison, WI, USA).

### Evaluation of antiviral efficacy of raloxifene

Cells were infected at 37°C for 1 h with the SARS-CoV-2 isolate at a MOI of 0.05. Infection was carryed out in DMEM without FBS. Then, the virus was removed and cells washed with warm phosphate buffered saline (PBS) and cultured with medium containing 2% FBS in the presence or in the absence of raloxifene. The compound was used at the concentration of 1.25, 2.5, 5, 10 and 15 µM and both cells and supernatants were collected for further analysis 48 h post infection (p.i).

### Plaque Assay

Cells were seeded at a density of 5×10^5^ cells/well in a 12-well plate and incubated at 37°C for 24 h. Supernatants from infected cells were serially diluted in DMEM without FBS and added to the cells. After 1 h incubation, media were removed and cells washed with warm PBS. Then cells were covered with an overlay consisting of DMEM with 0.4% SeaPlaque (Lonza, Basel, Switzerland). The plates were further incubated at 37°C for 48 h. Cells were fixed with 10% formaldehyde at room temperature for 3 h. Formaldehyde was aspirated and the agarose overlay was removed. Cells were then stained with crystal violet (1% CV w/v in a 20% ethanol solution), and viral titer (PFU/mL) of SARS-CoV-2 was determined by counting the number of plaques.

### Viral RNA extraction and quantitative real-time RT-PCR (qRT-PCR)

RNA was extracted from clarified cell culture supernatants (16,000 g x 10 min) and from infected cells using QIAamp Viral RNA Mini Kit and RNeasy Plus mini kit (Qiagen, Hilden, Germany), respectively, according to the manufacturer’s instructions.

RNA was eluted in 30 μl of RNase-free water and stored at -80 °C till use. The qRT-PCR was carried-out following previously described procedures with minor modifications [69]. Briefly, reverse transcription and amplification of the S gene were performed using the one-step QuantiFast Sybr Green RT-PCR mix (Qiagen) as follows: 50 °C for 10 min, 95 °C for 5 min; 95 °C for 10 sec, 60 °C for 30 sec (40 cycles) (primers: RBD-qF1: 5’-CAATGGTTTAACAGGCACAGG-3’ and RBD-qR1: 5’-CTCAAGTGTCTGTGGATCACG-3). A standard curve was generated by determination of copy numbers derived from serial dilutions (10^3^-10^9^ copies) of a pGEM T-easy vector (Promega, Madison, WI, USA) containing the receptor binding domain of the S gene (primers: RBD-F: 5’-GCTGGATCCCCTAATATTACAAACTTGTGCC-3’; RBD-R: 5’-TGCCTCGAGCTCAAGTGTCTGTGGATCAC-3’).

### Western blot analysis

Western blot was carried-out following previously described procedures with minor modifications [70]. Protein samples (30 µg) obtained from lysis in RIPA buffer (Cell Signaling Technology, Danvers, MA, USA) of infected cells were separated on 10% SDS-PAGE and then transferred onto polyvinylidene difluoride (PVDF) membranes (Millipore, Sigma, Burlington, MA, USA). After being blocked with 3% BSA in TBS buffer containing 0.05% Tween 20, the blot was probed with a human serum (1:1000 dilution) containing IgG to the SARS-CoV-2 nucleoprotein (NP) and with mouse anti-human GAPDH monoclonal antibody (G-9: Santa Cruz Biotechnology, Dallas, TX, USA). The antigen-antibody complexes were detected using peroxidase-conjugated goat anti-human or goat anti-mouse IgG (Sigma) and revealed using the enhanced chemiluminescence (ECL) system (Santa Cruz Biotechnology).

### Evaluation of antiviral efficacy of raloxifene on SARS-CoV2 variants

Vero E6 cells were infected at 37°C for 1 h with the SARS-CoV-2 strains indicated in the Virus section at a MOI of 0.05 in 96 well plates. Infection was carryed out in MEM (Sigma) without FBS (Gibco). Then, the virus was removed and cells washed with warm phosphate buffered saline (PBS) and cultured with medium containing 2% FBS in the presence or absence of raloxifene at different doses (0.23, 0.47, 0.94, 1.88, 3.75, 7.5, 15 µM) at 37°C and 5% CO2 up to 72 h. To determine antiviral efficacy of raloxifene, cell viability and viral induced CPE were mesured in not infected and infected cells treated with serial dilution of the drug, staining the cells with a solution of Crystal Violet (Diapath) and 2% formaldehyde. After 30 min, the fixing solution was removed by washing with tap water, and cell viability was measured by a photometer at 595 nm (Synergy™ HTX Multi-Mode Microplate Reader, Biotek, Winooski, VT, USA).

The percentage of viable cells for each condition was calculated compared to infected-not-treated (set as 0%) and not-infected-not-treated cells (set as 100%). The effect of raloxifene on cell viability was also checked by crystal violet/ 2% formaldheyde staining in each experiment performed for SARS-CoV-2 variants study.

### Data analysis

The half-cytotoxic concentration (CC_50_) and the half-maximal inhibitory concentration (IC_50_) for raloxifene were calculated from concentration-effect-curves after non-linear regression analysis using GraphPad Prism8. The selectivity index (SI) for raloxifene was calculated as the ratio of CC_50_ over IC_50_ [71].

### Statistical analysis

Data for the *in vitro* experiments performed were analyzed for statistical significance using the 1-way ANOVA, and the Bonferroni post-test was used to compare data. Differences were considered significant when *p* < 0.05. Statistical tests were performed using GraphPad Prism 8.

## Results

### In vitro effects of raloxifene on metabolism and SARS-CoV-2 infection in different cell lines

At first, a standard assay was carried out to measure the activity of raloxifene on cell metabolism. To this end, Vero E6 cells were cultured for 48 h in the absence or presence of different drug concentrations (range from 1.25 µM to 30 µM). As shown in Figure 1A, raloxifene-treated Vero E6 cells showed a normal surface-adherent phenotype until the concentration of 15 µM. A drug-dependent cytopathic effect was evident at concentration of 20 µM, involving the entire monolayer at a concentration of 25 µM and 30 µM. At the same time, raloxifene shows a slight effect on the extent of cellular ATP accumulation at a concentration ranging from 1.25 μM to 15 μM (87% and 70%, respectively). At higher doses, raloxifene showed a dose-dependent effect on ATP accumulation, reaching 56%, 35% and 0.6% at 20 µM, 25 µM and 30 µM (Figure 1B). The CC_50_ of raloxifene in Vero E6 cells was determined to be 18.4 µM.

**Figure 1.**
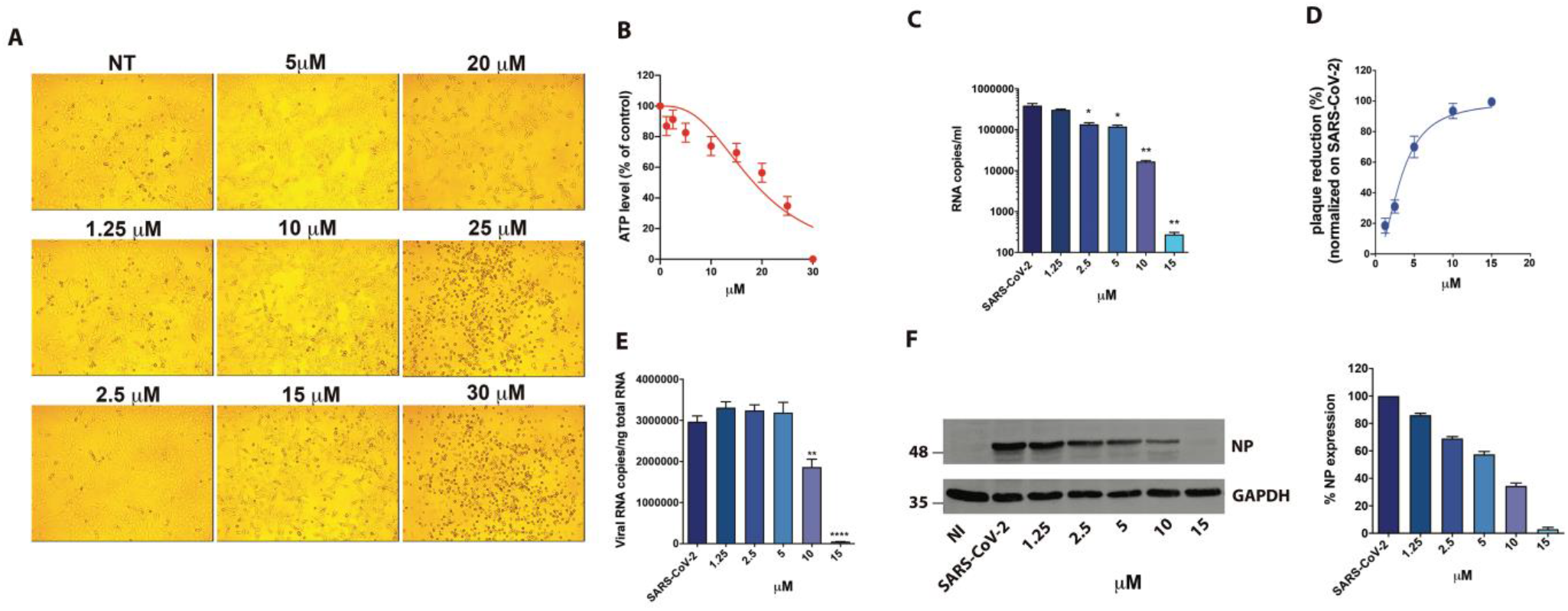
Effect of Raloxifene on Vero E6 cells. Vero E6 cells were cultured for 48 h in the absence or in the presence of raloxifene at different concentrations. (**A**) 10× bright-field images of Vero E6 cells after incubation for 48 h at 37°C with the indicated raloxifene concentrations. (**B**) CellTiter-Glo was used to measure the antimetabolic effect of raloxifene. (**C-F**) Vero E6 cells were infected with SARS-CoV-2 and cultured in the absence or in the presence of different doses of raloxifene. (**C**) Viral yield in cell supernatants was quantitated by qRT-PCR. (**D**) Viral titer in cell supernatants was evaluated by plaque assay and plotted as percentage of plaque reduction compared to SARS-CoV-2. (**E**) Quantitation of SARS-CoV-2 genomes at the intracellular level by qRT-PCR. (**F**) NP expression in infected cells was analyzed by western blot (left panel). Densitometric analysis of western blot is shown in the right panel. Graph represents the percentage of NP expression. Data are representative of two independent experiments with similar results. All the experiments were performed at least in three independent replicates and pictures shown are representative. Data are presented as the mean ± standard error of the mean *, P < 0.05; **, P < 0.01; ****, P < 0.0001.

Next, to assess the antiviral activity of raloxifene, Vero E6 cells were infected with SARS-CoV-2 (B.1 lineage) at a MOI of 0.05 [67]. Specifically, Vero E6 were infected with SARS-CoV-2, and 1 h later cultured in the absence or presence of different raloxifene concentrations (range from 1.25 µM to 15 µM). Raloxifene efficiently inhibits viral replication. In particular, viral genome copy numbers evaluated on supernatants collected at 48 h p.i. by qRT-PCR, showed a significant reduction of the virus yield already at 2.5 µM raloxifene concentration (2.9-fold reduction), with a maximal reduction at 15 µM (1400-fold reduction) (Figure 1C). Raloxifene also displayed a dose-dependent inhibition of viral replication in Vero E6 cells, as determined by infectious viral titers, exhibiting a 70% reduction of viral titer at a concentration of 5 µM, with 94% to 100% inhibition at 10 µM and 15 µM, respectively (Figure 1D). Raloxifene efficacy was then confirmed at intracellular level. Quantification of viral RNA in SARS-CoV-2-infected cells showed a significant reduction of intracellular SARS-CoV-2 genome copy number already at 10 µM and a 99-fold reduction at 15 µM (Figure 1E).

Accordingly, western blot (WB) analysis showed a dose-dependent inhibition of SARS-CoV-2 upon raloxifene treatment with 43% reduction of NP viral protein expression at a concentration of 5 µM, with 65% and 97% reduction at 10 µM and 15 µM, respectively (Figure 1F). Raloxifene IC_50_ value was calculated to be 3.3 µM. SI was then calculated and found to be 5.6. We then tested the raloxifene activity on Calu-3 cells as a model of human pulmonary cell line. The CC_50_ value was determined, as above, and found to be 24.4 µM (Figure 2A). Next, Calu-3 cells were infected as described above. Supernatants were collected at 48 h p.i., and tested for viral genome copy numbers by qRT-PCR. As shown in Figure 2B, the treatment significantly reduced the virus yield. In particular, raloxifene displayed a dose-dependent inhibition of viral replication, as determined by infectious viral titers, exhibiting a 67% reduction of viral titer at a concentration as low as 10 µM, with 96% and 98% inhibition at drug concentrations of 15 µM and 25 µM, respectively. The efficacy of the treatment was confirmed at intracellular level by qRT-PCR and WB on NP (Figures 2C-E). The IC_50_ was calculated and found to be 9 µM. SI was then calculated and found to be 2.7.

**Figure 2.**
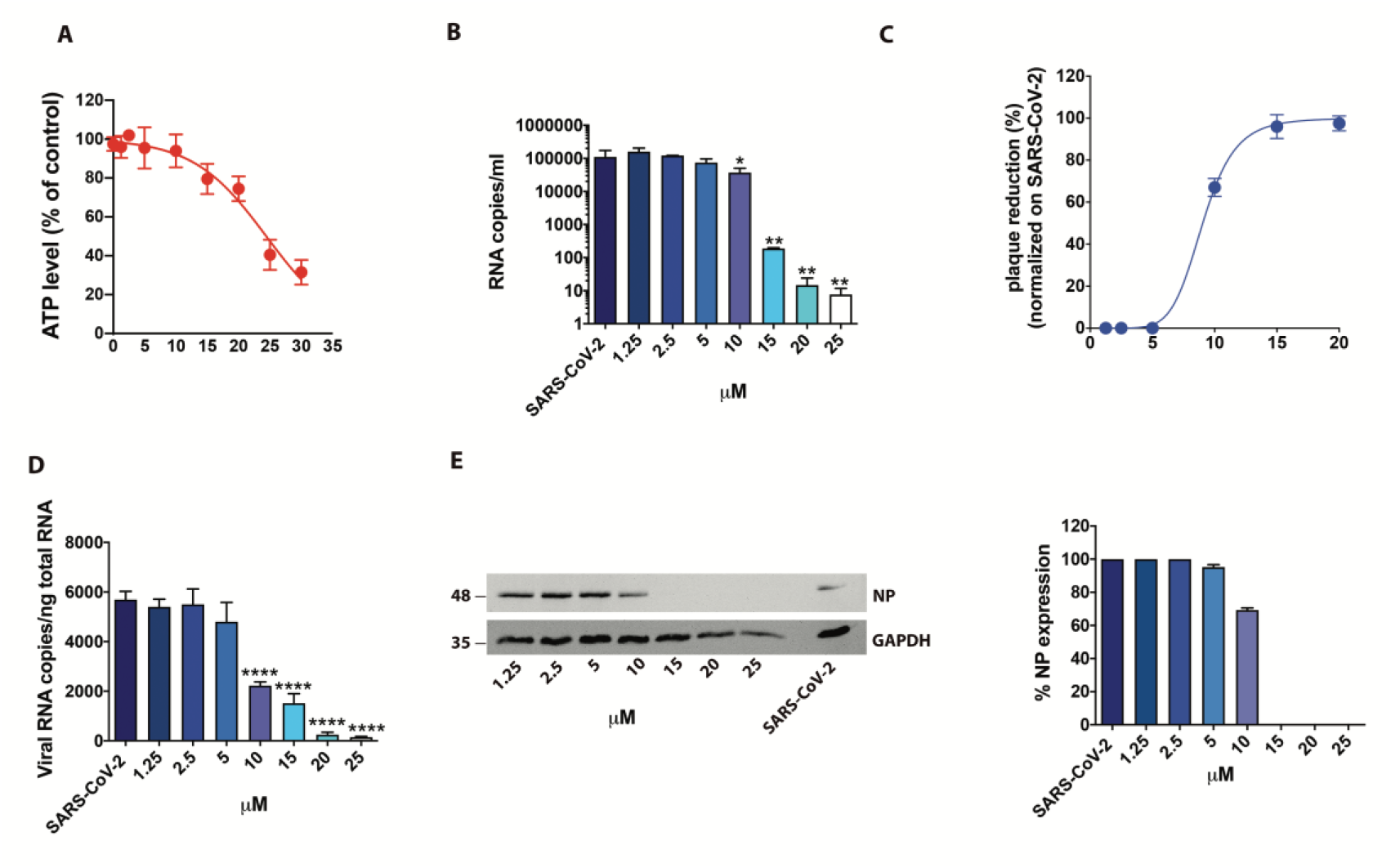
Effect of Raloxifene on Calu-3 cells. **(A)** Calu-3 cells were cultured for 48 h in absence or in the presence of raloxifene at different concentrations. CellTiter-Glo was used to measure, the antimetabolic effect of raloxifene. (**B-E**) Cells were infected with SARS-CoV-2 and cultured in the absence or in the presence of different doses of raloxifene. (**B**) Viral yield in cell supernatants was quantitated by qRT-PCR. (**C**) Viral titer in cell supernatants was evaluated by plaque assay and plotted as percentage of plaque reduction compared to SARS-CoV-2. (**D**) Quantitation of SARS-CoV-2 genomes at the intracellular level by qRT-PCR. (**E**) Nucleocapsid (NP) protein expression in infected cells was analyzed by western blot (left panel). Densitometric analyses of western blot results are shown. Graph represents the percentage of NP protein expression. Data are representative of two independent experiments with similar results. All the experiments were performed at least in three independent replicates and pictures shown are representative. Data are presented as the mean + standard error of the mean *, P < 0.05; **, P < 0.01; ****, P < 0.0001.

### Raloxifene exerts antiviral activity on SARS-CoV-2 variants

We then performed a systematic study of the antiviral efficacy of raloxifene on the most common variants on Vero E6 cells (Figure 3). Different viral strains were used: the wild type isolated in January 2020 from Chinese patient (named Wuhan), two different isolates for the D614G spike variants representing the dominant strains circulating in Europe from April to December 2020 (named GV and D614G) and the variants of major concern originated in UK, Brazil, South Africa and India (named VOC B1.1.7, VOC P.1, VOC B1.351, and VOC B1.617.2, respectively). We first determined the time-window in which CPE appeared for each variant. The CPE was evident at 48h for all the tested strains but VOC B1.1.7 and VOC P.1 variant, for which evident CPE appeared later (56h and 72h, respectively). In parallel, uninfected cells were cultured in the presence of different doses of raloxifene to evaluate possible cytotoxicity due to the treatment. In cells treated with drug at 15 µM, we observed a reduced percentage of viable cells as revealed by crystal violet staining (82.9 +/-8.69% and 76.3+/-5.84%, at 48 and 72h respectively, compared to 100% in untreated cells). No significant effect on cell viability with lower dug concentrations (7.5 to 0.23 µM) was observed. To determine antiviral efficacy of the drug, CPE was measured in infected cells treated with seven two-fold serial dilutions of raloxifene (15 to 0.23 µM) using the time windows identified for each strain. The drug was able to recover cell viability in Vero E6 cells infected with all the tested viral strains. The half-maximal inhibitory concentration (IC_50_) calculated on recovering of cell viability varied from 4.50 to 7.99 µM depending on the strain (Figure 3), showing a strong antiviral activity against all the variants under investigation.

**Figure 3.**
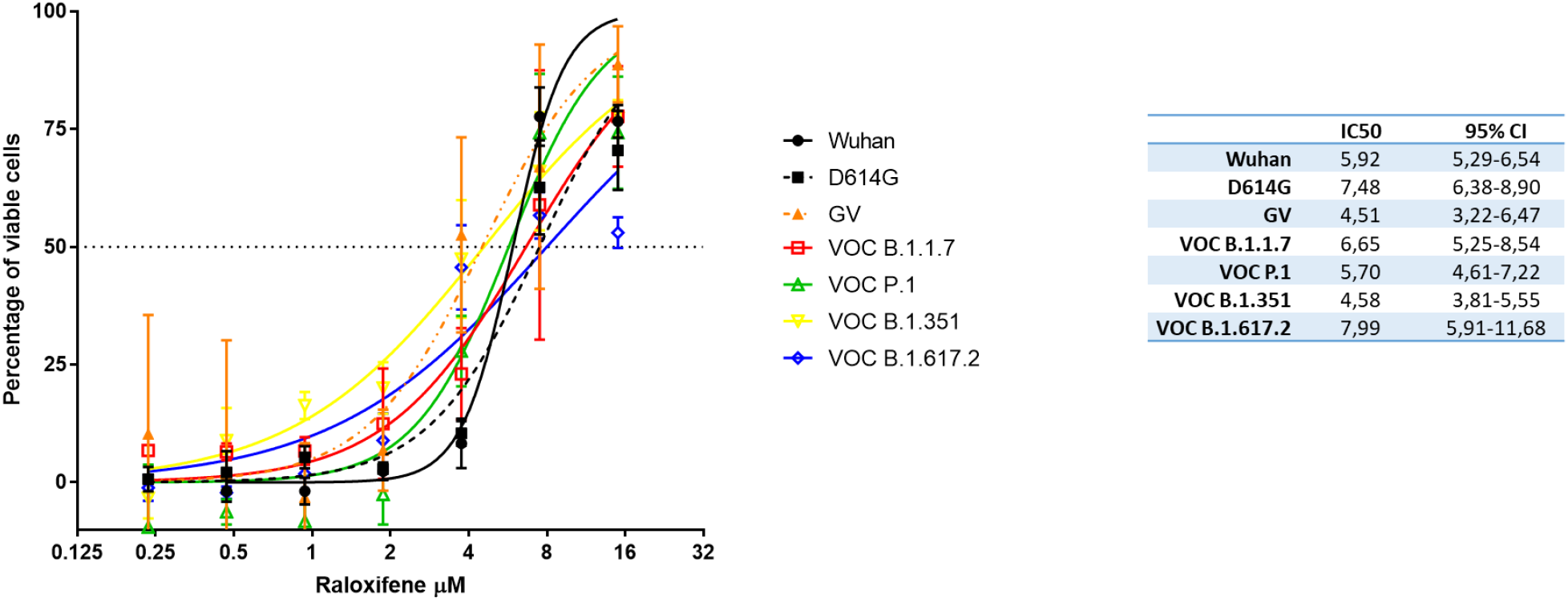
Raloxifene reduces the cytopathic effect (CPE) induced by SARS-CoV-2 variants in Vero E6 cells. The graph shows the inhibition of CPE observed at different concentration of raloxifene. The IC50 calculated by non linear regression are shown in the table. Percentage of viable cells calculated on not treated not infected = 100%; not treated SARS-CoV-2 infected cells= 0%. Bars indicate SD.

### System Biology screening to investigate poly-pharmacological effects of raloxifene against SARS-CoV-2 infection

Besides being able to directly bind viral proteins such as Spike, raloxifene may exert a range of other modulatory effects on the infection by SARS-CoV-2. In a recent paper we reviewed the possible links between ER modulation and host response to viral infections against different viruses, suggesting a therapeutic potential for SERMs in the control of COVID-19 infection [38]. Aiming to strengthen the above hypothesis we built a molecular network connecting the human-virus interactome and those proteins known to be involved in the COVID-19 pathogenesis, as described in the Materials & methods section. The resulting network was in turn used to generate a list of proteins; each member of this list was used as a probe to screen all the papers on raloxifene nominating the proteins relevant for the SARS-CoV-2 infection. In this way, three functional groups of human genes involved in the biology of the viral infection and potentially modulated by raloxifene were identified:

1. A group of genes including those modulated by the raloxifene molecular target, specifically ESR2, and connected to inflammation;
2. A group of genes including those expressed in the lungs which are modulated by the raloxifene molecular target, specifically ESR; when deregulated, the corresponding genes cause severe asthma, in agreement with the enrichment in this group of genes whose unfavorable variants cause worse respiratory consequences, according to GWAS studies;
3. A group of genes directly modulated by the virus, both during the cell entry phase and the replication phase, which also include proteins upstream or downstream of some raloxifene-controlled pathways.

Taken all together, these results suggest a potential poly-pharmacological effect of raloxifene in COVID-19 as anti-inflammatory, respiratory and antiviral.

### Effect on inflammation

Looking at the genes identified as linked to the promotion of inflammation by the virus, it is relevant to keep in mind that one of the clinically validated targets of the anti COVID-19 therapy is the cytokine Interleukin-6 (IL-6) [72]. A downregulation of the inflammatory signal and of the IL-6 expression was found with raloxifene in a clinical setting [73]. Besides IL-6, other serum cytokines (i.e. TNF-alpha and TGF-beta1) involved in the cytokines storm due to SARS-CoV-2 are regulated by raloxifene.

### Effect on respiratory response

Genes regulating the production of nitric oxide are implicated in the vascular and respiratory response to the viral infection. Treatment of rats with raloxifene was shown to upregulate the expression of eNOS (NOS3) in rat thoracic aorta, after complexing with the ESR_2_ expressed in lungs. This is expected to exert a potentially important vasculo-protective effect, and eventually to contribute to clinical improvements in ARDS and pulmonary hypertension [74]. Other compounds, e.g. Rho kinase (ROK) inhibitors, phosphodiesterase-5 inhibitors [75], omentin [76] were also shown to activate the eNOS (NOS3) pathway, with protective effect for ARDS and related inflammation in experimental models. In ARDS patients, the effects of inhaled nitric oxide on the reduction of pulmonary blood pressure and on the improvement of oxygenation, offered the rationale for a clinical trial in severe COVID-19 patients (NCT04388683).

### Effect on antiviral actio

Consistently with the identification of the genes from the GWAS study, a direct antiviral action of raloxifene, in terms of inhibition of viral replication and/or infection, was found in several different contexts like in vitro against EbolaVirus [58, 59, 77], against HBV [62], and against HCV [60]; in human female cells from nasal epithelium against the Influenza Virus A [64], and also in a randomized clinical trial on 123 postmenopausal women, against HCV [65].

## Discussion

Raloxifene, a second generation SERM, was previously proposed as a potential candidate for the treatment of COVID-19 patients due to the *in silico* predicted possibility to interfere with the viral replication and disease progression with multiple mechanisms of action both ER dependent and independent [38].

In this paper we report for the first time the *in vitro* characterization of the antiviral activity of raloxifene against SARS-CoV-2 infection using two relevant experimental systems, Vero E6 monkey kidney cells and human pulmonary Calu-3 cells. SARS-CoV-2 infected monkey Vero E6 cells are commonly used to study coronavirus infection as they support viral replication to high titres and highly express ACE-2 receptor [78-82] that plays an essential role for SARS-CoV-2 entry into the cells [83]. SARS-CoV-2 infected human Calu-3 cells are a relevant and predictive model because of airway epithelial origin [84]. In both assays the results confirmed that raloxifene blocks with high efficiency SARS-CoV-2 replication.

The characterization was completed testing raloxifene also against all the most common circulating SARS-CoV-2 variants of clinical relevance, confirming that it maintains a high and consistent activity, thus reinforcing the interest on its potential clinical use as antiviral agent in COVID-19 patients.

Raloxifene cytotoxicity was assessed with two independent assays: the first measuring the activity of the compound on cell replication, the second on cellular metabolism. With both approaches we found that the CC50 was attested at high micromolar range, which is far from the low micromolar range in which the antiviral activity was observed. The selectivity index (SI) of the drug was superimposable in the two experimental models in the range of 2 to 7. In general, the value of SI for a drug with direct antiviral activity is greater than 1; the higher the SI value, the more effective and safer the drug is. Some authors [85-87] report a limit value of SI = 4 to define a compound as a good compound with direct antiviral activity. The SI value found in the models is therefore indicative of a molecule with a significant antiviral activity and with an activity/toxicity profile consistent with a possible translation to human clinical trials. In addition, raloxifene is a drug that has been used for a long time, and its safety profile is supported by a huge volume of clinical data from long term treatments [88-90]. The occurrence of thromboembolic events, even though rare, in patients treated with raloxifene has to be regarded with particular caution due to the high risk of thromboembolic manifestations in COVID patients. A short duration of treatment and the careful avoidance to treat patients with concomitant risks of thromboembolic events are recommended.

Among SERMs raloxifene has a unique risk/benefit profile built on a large safety database not limited to oncological patients, like for other SERMs, but on a large use in postmenopausal women for the management of osteoporosis, including men treated for a variety of indications. The potential of SERMs, and in particular of raloxifene, found a promising confirmation in a recent retrospective study on a large population of cancer patients that demonstrated a protective effect on SARS-CoV-2 infection and a significant reduction of severity and duration in the subpopulation of patients treated with raloxifene [91].

A system biology study was also conducted with the aim to match the available information on gene and pathways regulated by raloxifene against a Cytoskape-generated human SARS-CoV-2 interactome network. The results of the study strongly support the concept that raloxifene may positively influence the course of SARS-COV-2 infection by modulating three functional groups of human genes, all of them playing a key role in the biology of the viral infection. The systematic data analysis went further, confirming the putative antiviral activity, and also highlighting the potential of raloxifene to exert both an antinflammatory action by downregulating the expression of key mediators of the cytokine storm, and a vasculo-protective effect by upregulating eNOS expression (NOS3) [74]. These findings on one hand are in agreement with previous papers highlighting the protective effect of estrogen signalling in the context of COVID-19 infection, and on the other hand confirm the specific characteristics of raloxifene as an ideal candidate to put to a test the hypothesis in the clinics due to ist peculiar mechanism within the class of SERMs and its potential ability to exert a pleiotropic effect by targeting viral and host targets with a key role in the disease progression and exacerbation.

